# Deletion of the creatine transporter gene in neonatal, but not adult, mice lead to cognitive deficits

**DOI:** 10.1101/582320

**Authors:** Kenea C. Udobi, Nicholas Delcimmuto, Amanda N. Kokenge, Zuhair I. Abdulla, Marla K. Perna, Matthew R. Skelton

**Affiliations:** Department of Pediatrics, University of Cincinnati College of Medicine and Division of Neurology, Cincinnati Children’s Research Foundation. Cincinnati, OH 45229

## Abstract

Creatine (Cr) is a guanidino compound that provides readily-available phosphate pools for the regeneration of spent ATP. The lack of brain Cr causes moderate to severe intellectual disability, language impairment, and epilepsy. The most prevalent cause of Cr deficiency are mutations in the X-linked *SLC6A8* (Creatine transporter; CrT) gene, known as CrT deficiency (CTD). There are no current treatments for CTD and the mechanisms that underlie the cognitive deficits are poorly understood. One of the most critical areas that need to be addressed is if Cr is necessary for brain development. To address this concern, the *Slc6a8* gene was knocked out in either neonatal (postnatal day (P)5) or adult (P60) mice. The P5 knockout mice showed deficits in the Morris water maze and novel object recognition, while there were no deficits in P60 knockout mice. Interestingly, the P5 knockout mice showed hyperactivity during the dark phase; however, when examining control mice, the effect was due to the administration of tamoxifen from P5-10. Taken together, the results of this study show that Cr is necessary during periods of brain development involved in spatial and object learning. This study also highlights the continued importance of using proper control groups for behavioral testing.

**Take-home message:** The learning and memory deficits seen in Slc6a8-deficient mice are likely due to the developmental loss of Cr.

## INTRODUCTION

Creatine (Cr) is a guandino compound that serves as a phosphate reserve to replenish spent ATP (Wyss and Kaddurah-Daouk 2000). At the mitochondria, Cr kinase (CK) transfers a phosphate from ATP to Cr, creating phospho-Cr (pCr). At sites of energy utilization, CK transfers the phosphate from pCr to ADP, thereby regenerating ATP. As Cr is significantly smaller than ATP, these pCr pools allow for a greater energy density in cells. The importance of Cr to cellular function is highlighted by a group of disorders resulting in the lack of brain Cr (Schulze 2003). Caused by a lack of Cr synthesis or transport, cerebral Cr deficiency leads to intellectual disability, a lack of language development, and epilepsy. The most common cause of Cr deficiency is due the loss of the *SLC6A8* gene (Cr transporter; Crt)(Cecil et al 2001; deGrauw et al 2003). Crt deficiency (CTD) is one of the most common causes of X-linked intellectual disability, with an estimated carrier frequency of 1 in 4,200 (DesRoches et al 2015). Unlike deficits in Cr synthesis, CTD cannot be treated with oral Cr supplementation (van de Kamp et al 2013) or Cr synthesis precursors (Valayannopoulos et al 2012; Dunbar et al 2014; Bruun et al 2018). The inability to restore Cr in CTD makes the development of molecules that could deliver Cr past the blood-brain barrier (Trotier-Faurion et al 2013) or Cr mimetics (Kurosawa et al 2012) valuable as possible treatment strategies. However, the lack of understanding on the role of brain Cr may lead to an inadequate design for proof of concept studies.

One of the primary concerns in designing treatment protocols should be targeting the critical periods in which Cr is necessary to facilitate cognitive processes. It has been shown that early-life Cr supplementation provides a greater cognitive improvement than later life treatment in individuals with mutations of the Cr-synthesis genes *Arginine:glycine amidinotransferase* (*AGAT*) or *guanidino-methyltransferase (GAMT)* (Battini et al 2006; Stockler-Ipsiroglu et al 2014; Stockler-Ipsiroglu et al 2015). Mitochondrial dysfunction or CK inhibition reduces the growth and dendritic arborization of cerebellar Purkinje cells *in vitro* (Fukumitsu et al 2015). The neurons with dysfunctional mitochondria were rescued by Cr supplementation, highlighting the role Cr plays in cellular metabolism. Together, the human and *in vitro* data strongly suggest that Cr is necessary for brain development.

We developed mice that have exons 2-4 of the *Slc6a8* gene flanked by loxP sites (*Slc6a8*^*FLOX*^). Ubiquitous (*Slc6a8*^*−/y*^) and brain-specific knockout mice generated from this line have spatial learning and memory, novel object memory, and conditioned fear deficits (Skelton et al 2011; Udobi et al 2018). This wide range of deficits mirror the global cognitive delay seen in CTD patients. The purpose of this study was to determine if the neonatal or adult loss of the *Slc6a8* gene was sufficient to cause learning and memory deficits. Using a tamoxifen-inducible CreER^T2^, we examined the cognitive effects caused by deletion of the *Slc6a8* gene on postnatal day (P)5 or P60.

## METHODS

Full methods are available in the supplement.

### Generation of conditional knockout mice

All institutional and national guidelines for the care and use of laboratory animals were followed. Female *Slc6a8*^FLOX/+^ mice, generated and genotyped as described (Skelton et al 2011), were mated to mice expressing an inducible Cre recombinase driven by the *UBC* promoter (B6.Cg-*Ndor*^*Tg(UBC-Cre/ERT2)Ejb*^*/1J (UBC-CreERT2)*; Jackson laboratory, Bar Harbor, ME). On either P5 or P60, mice were treated (1/d i.p.) with tamoxifen (75 mg/kg) or corn oil vehicle for 5 d, creating neonatal (P5KO) or adult (P60KO) mice. Behavioral testing was performed 60 d following treatment. The groups (along with N’s) used are outlined in supplemental table 1.

### *Slc6a8* expression and Cr determination

#### Tissue collection

Mice were lightly anesthetized with isoflurane inhalation and sacrificed by decapitation. The brain was removed from the skull, divided into hemispheres, and flash frozen.

#### Cr determination

Cr content was measured using reversed phase HPLC with UV detection (Tranberg et al 2005).

#### Quantitative RT-PCR

Real-time quantitative PCR (QPCR) was used to measure *Slc6a8* expression levelsin the brain as described previously (Hautman et al 2014).

### Morris water maze

The Morris water maze (MWM) is a test of spatial learning and reference memory (Vorhees and Williams 2006). Mice were tested as described (Skelton et al 2011; Udobi et al 2018). Mice were tested in three phases, a visible platform phase, a hidden platform acquisition phase and a reversal phase. Hidden platform testing was conducted over 4 days with a probe trial on day 5 of testing. Performance was measured using ANY-maze® software (Stoelting Company, Wood Dale, IL).

### Novel object recognition

Novel object recognition (NOR) is a test of incidental learning and memory (Clark et al 2000). Mice were tested in the ANY-box apparatus (Stoelting Company, Wood Dale, IL) as previously described (Hautman et al 2014; Udobi et al 2018). On the test day, mice were presented with two identical objects and allow to explore until 30 s of observation time between objects was accrued. The maximum time for the trial was 10 min. One hour later, memory was tested by presenting the mouse with an identical copy of one of the familiar objects along with a novel object. Percent time spent observing the novel object was the independent variable for this test.

### Contextual and cued fear conditioning

Conditioned and contextual fear was assessed as described with modification (Peters et al 2010; Udobi et al 2018).

### Overnight locomotor activity

Spontaneous locomotor activity (Brooks and Dunnett 2009) was tested in automated activity chambers. Mice were placed into the open-field arena at 1800 and left undisturbed for 14 h, with lights off at 2000 and on at 0600 the next day. The dependent measure was total number of photobeams interruptions.

### Statistics

The hypothesis of this study was that the conditional knockout mice would differ from corn-oil treated *Slc6a8*^*+/y*^*::Ubc-CreERT2*^*−*^ mice (WT-VEH). Therefore, preplanned comparisons were designed regardless of the outcome of the omnibus ANOVA. As there were no interactions with the repeated measure, the Dunnett’s tests were performed using the means of all trials. A p<0.05 is used to reject the null hypothesis that the groups are the same. The F values from the omnibus ANOVAs for each test are presented in Supplemental Data Table 2. Data are presented as LSMEANS±SEM and will be made freely available upon request.

## RESULTS

### Body weight, *Slc6a8* expression, Cr measurement

The expression of *Slc6a8* mRNA was not observed 5 days following tamoxifen administration in either the P5KO or the P60KO mice (data not shown). Brain Cr levels were reduced in both P5KO (Figure 1A) and P60KO (Figure 1B) mice. P5KO (Figure 1C) and P60KO (Figure 1D) weighed less than the control mice at the time of behavioral testing. No effects of tamoxifen were observed. We have shown that these body weight reductions do not affect cognitive function (Udobi et al 2018).

**Figure 1.**
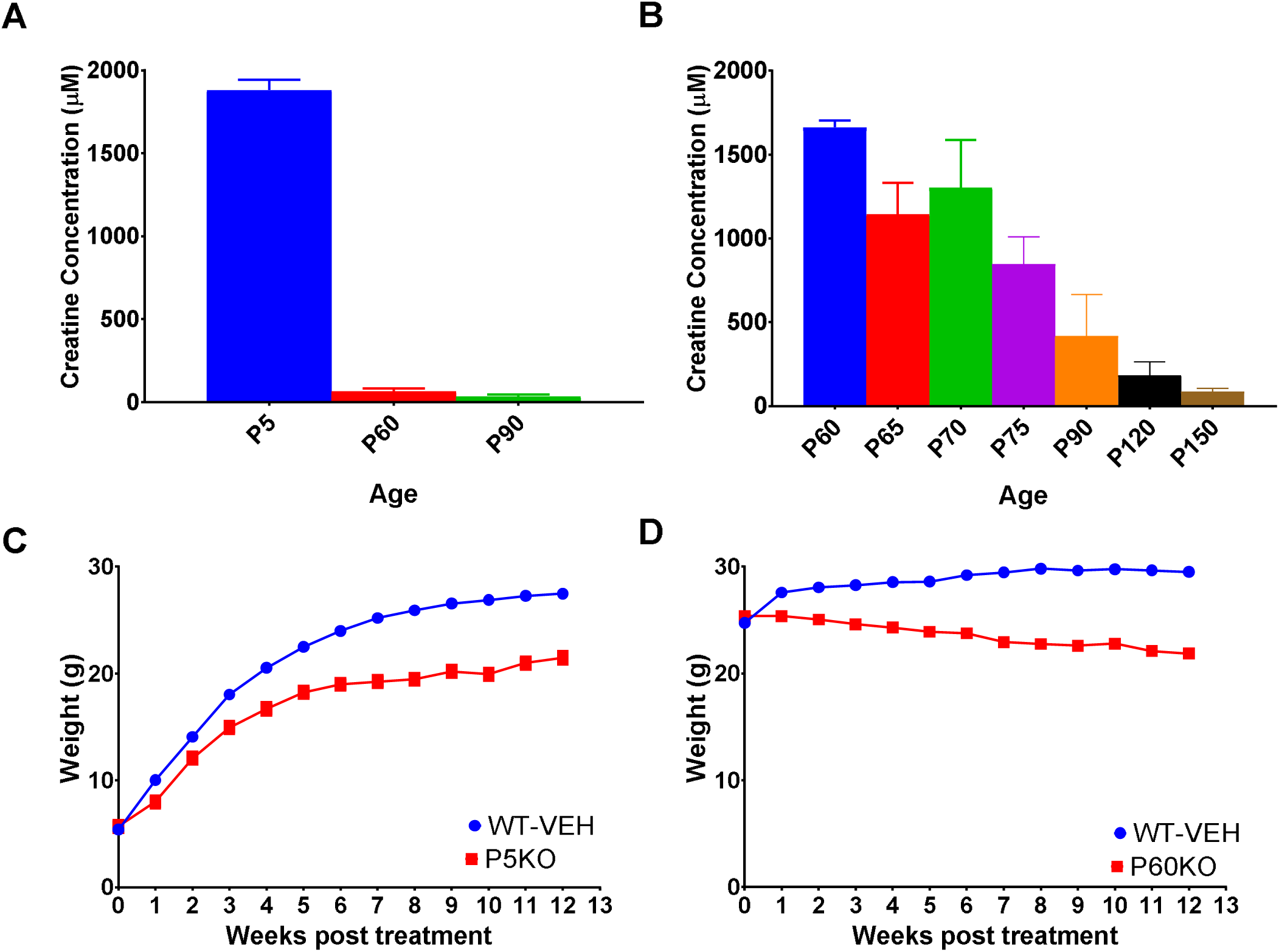
Reductions in body weight and brain Cr levels following P5 or P60 *Slc6a8* deletion. A) Brain Cr levels in P5KO mice before and 60- or 90-days following tamoxifen exposure. By the time of behavioral testing (P60) Cr levels were reduced to less than 10% of initial levels. B) Time course of Cr levels in P60KO mice. C) The P5KO mice gained weight as they aged however the growth did not match WT mice (main effect of group). D) In P60KO mice there was a body mass reduction following tamoxifen administration. Data are LSMEAN±SEM. N=13-18/group

### Morris Water Maze

On the first day of visible platform testing, P5KO and WT-VEH mice had a similar latency to the platform (t_130_=2.23, p=0.1386, Figure 2A). On days 2 and 3, P5KO mice had longer latencies across days compared with the WT-VEH group (t_130_=3.03, p=0.0181). For the P60-treated group, no effects were seen between P60KO and WT-VEH mice on any day (t_60.9_=2.04 p=0.2369 for days 2-3; Figure 2B).

**Figure 2.**
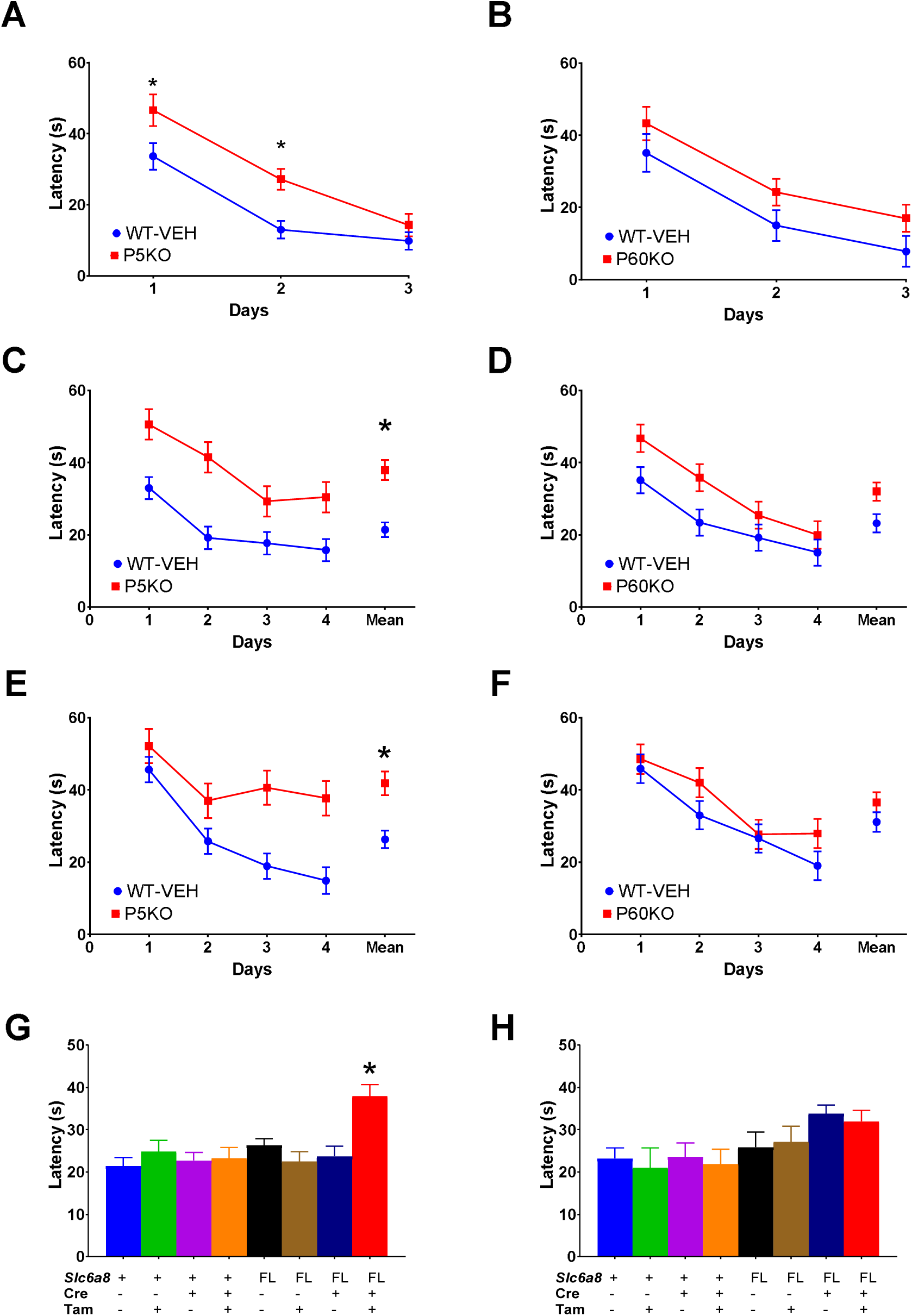
Spatial learning deficits in P5KO but not P60KO mice. During visible platform training (A), P5KO mice show an increase latency on days 1 and 2 compared with WT-VEH mice. All mice improved across days. In the hidden platform training, the P5KO mice show higher mean latencies to the platform across all days (Mean) during the (B) acquisition and (C) reversal phases of the MWM compared with the WT-VEH mice. (D) Shows the overall mean across days for all groups tested with only the P5KO mice differing from the WT-VEH mice. No differences were observed in the P60 during (E) visible platform, (D) acquisition, or (E) reversal testing. There were no differences between any P60 group tested and WT-VEH during the acquisition phase (H). Data are LSMEAN±SEM, *p<0.05. N=13-18/group

During the acquisition phase of hidden platform testing, P5KO mice had longer latencies to find the platform than the WT-VEH mice (t_174_=4.27, p*=*0.0002; Figure 2C). Similar effects were seen in the distance taken to the platform and for path efficiency, which measures how close to a direct path the mouse takes to find the platform. No differences were observed between WT-VEH mice and the other control groups in any measure (Figure 2G). There were no differences in swim speed between groups. During the reversal phase, the P5KO mice had longer latencies than WT-VEH group (t_140_=− 2.78, p=0.033; Figure 2E) while no differences were observed between control groups.

No differences were observed during the acquisition or reversal phase in P60KO mice (Figures 2D, 2F, and 2H). All groups had a similar swim speed in the P60 groups. (t_97.5_=−2.17, p=0.1727 P60KO vs WT-VEH).

A probe trial was conducted the day following the completion of each phase of MWM training as a test of spatial memory. (Supplemental Figure 1). There were no differences observed in the P5 or P60 groups in the acquisition phase. During the reversal phase, the P5KO had a greater average distance from the platform compared with WT-VEH (t_137_=2.74, p=0.0408). No differences were observed in P60KO mice (t_77_=0.78, p=0.4353).

### Novel object recognition

The P5KO mice spent less time with the novel object compared with WT-VEH (t_113_=−2.78, p=0.0384; Figure 3). No differences were observed in the P60 group or any of control groups from the P5 or P60 treatment.

**Figure 3.**
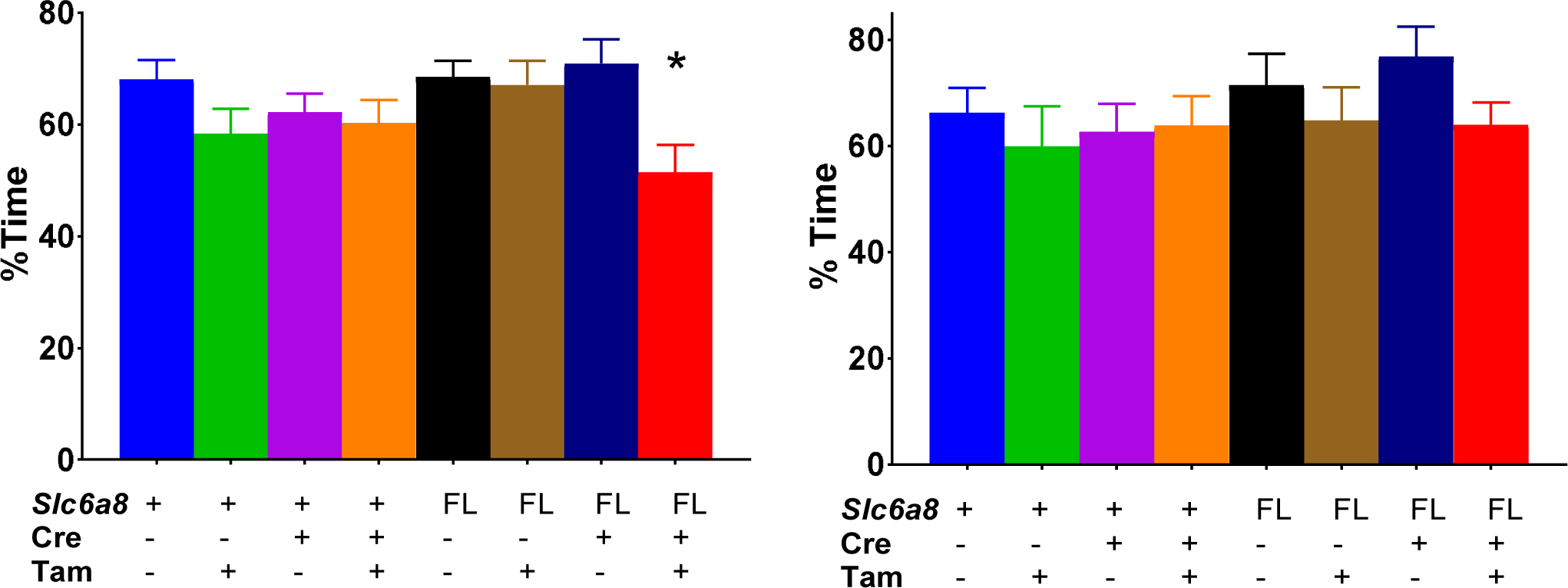
Novel object recognition deficits in P5KO mice. *Left:* In the P5-treated group, only the P5KO mice show a reduction in time spent with the novel object compared with the WT-VEH. *Right:* No differences from WT-VEH were observed in any of the groups treated at P60. Data are LSMEAN±SEM. N=9-18/group. *p<0.05

### Fear memory

No differences were observed in contextual or conditioned fear in either group (Supplemental Figure 2).

### Overnight locomotor activy

Locomotor activity was measured continuously for 2 h prior to the dark phase, the entire 10 h dark phase, and 2 h after the lights came back on (Figure 4). During the first 2 h, no differences were observed in either KO group. While the P5KO mice were hyperactive during the dark phase (t_459_=4.26, p=0.0002) there were significant effects of all tamoxifen controls vs WT-VEH. During the overnight testing, the tamoxifen-treated (t_211_=4.89. p<0.0001) and P5KO (t_211_=−2.96, p=0.0034) mice were hyperactive compared with vehicle-treated mice but did not differ from each other. In the 2 h light period following the dark cycle, tamoxifen-treated (t_144_=4.01. p<0.0001) and P5KO (t_144_=−2.26, p=0.0253) mice showed hyperactivity. No differences were observed between tamoxifen-treated and P5KO mice (t_211_=−0.31, p=0.7587). No differences in activity between the WT-VEH and the P60KO mice were observed during any phase.

**Figure 4.**
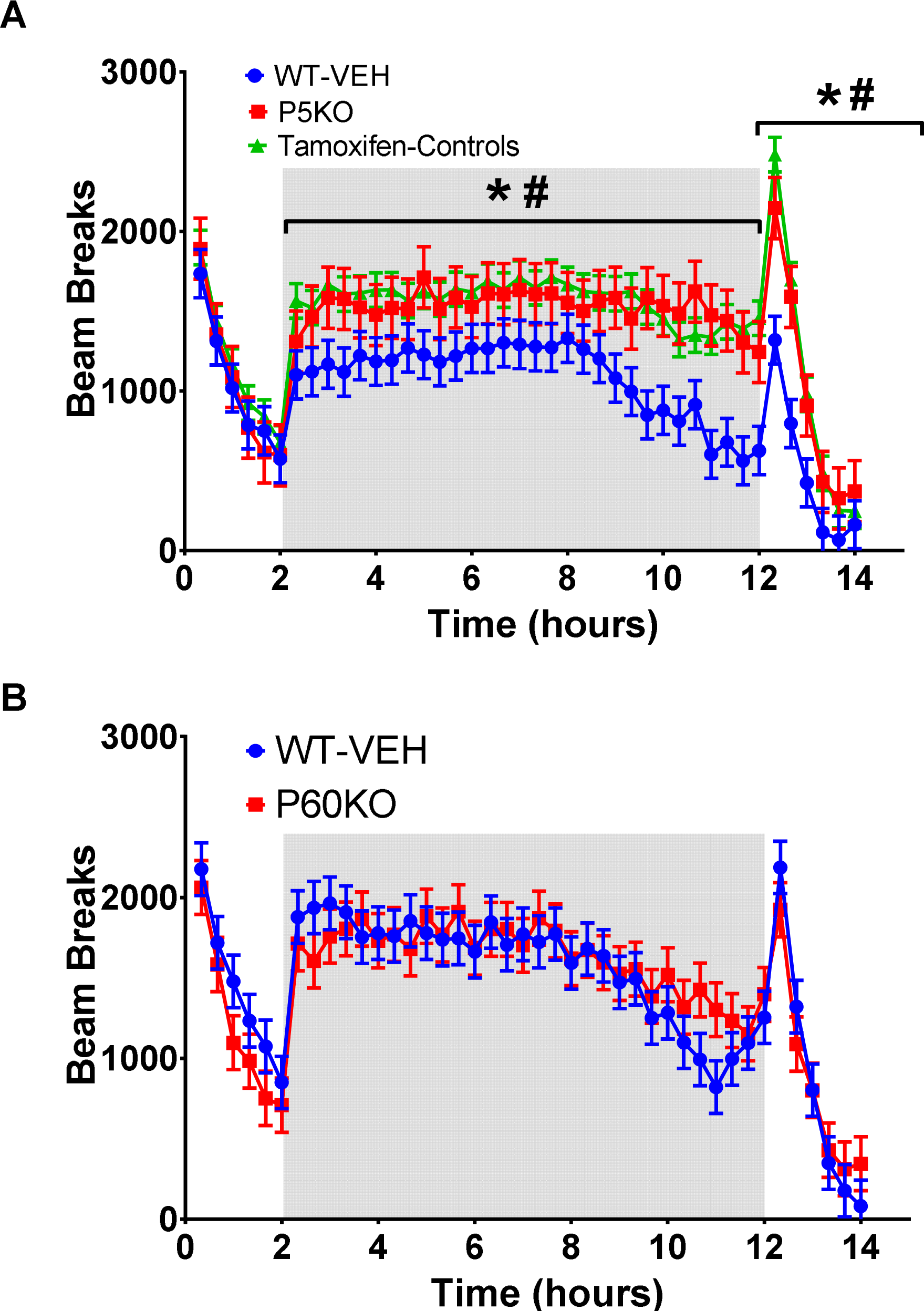
Neonatal tamoxifen administration leads to hyperactivity during adulthood. Locomotor activity was assessed in mice for 14 hours with the shaded region representing testing during the lights-off phase. Data are presented as the total beam breaks during a 20-min period. (A) The P5KO mice show a hyperactivity compared with WT-VEH mice; however, the remaining tamoxifen treated mice show a similar hyperactivity. (B) No differences in activity were observed between WT-VEH and P60K0. Data are LSMEAN±SEM. N=13-18/group. *p<0.05 P5KO vs WT-VEH; # p<0.05 Tamoxifen controls vs WT-VEH

### Stimulated respiration in synaptosomes

No differences were observed between P5 or P60 groups in stimulated respiration (Supplemental Figure 3).

## DISCUSSION

The purpose of this study was to determine if there is a developmental component to the cognitive deficits seen in CTD patients and *Slc6a8* knockout mice. By eliminating the *Slc6a8* gene in adulthood and during the perinatal period, we were able to determine that the later-developing brain is still sensitive to the loss of Cr while the adult brain is not. Combined with data from human Cr-synthesis deficiencies supplemented with Cr and in vitro work showing Cr is involved in neuronal morphology, these findings suggest that Cr plays an important role in brain development. In addition, we show that tamoxifen exposure during the perinatal period may lead to changes in locomotor activity. This finding further affirms the need for adequate controls in behavioral studies.

As seen in Figure 1, tamoxifen induced a successful recombination in *Slc6a8*^*FLOX*^ with no brain *Slc6a8* expression after 5 d of treatment. By the time of behavioral testing both groups of mice had Cr levels that were similar to Cr levels seen in the ubiquitous *Slc6a8* knockout mice (Skelton et al 2011). In the P60 mice, a gradual reduction in brain Cr levels was observed suggesting that Slc6a8 is necessary for the maintenance of brain Cr levels and not for the normal loss of Cr through degradation to creatinine. This is important when designing treatments for CTD as it suggests that there is not a “trapping” mechanism that could lead to an excessive amount of Cr accumulating with Cr-derived treatments. The loss of *Slc6a8* in adult mice caused an almost immediate weight reduction. The musculoskeleltal effects of Cr are well documented since most Cr-related studies are focused on its use as an athletic supplement. Slc6a8^−/y^ mice have an increased body fat percentage compared with WT mice (Perna et al 2016). Ubiquitous *Agat*^*−/−*^ mice are resistant to weight gain on a high-fat diet (Choe et al 2013), while mice with a fat-specific deletion of *Agat* weighed more than WT mice when fed a high-fat diet (Kazak et al 2017). Together, these data suggest that Cr may play an important role in obesity.

Translating rodent to human brain development is a multifactorial process without direct correlations in many cases. Many processes that happen simultaneously in the rodent do not occur simultaneously in humans. In the case of this study, we set out to understand why the lack of Cr leads to deficits in MWM, NOR, and conditioned fear performance. As MWM and NOR are largely mediated by the hippocampus, we concluded that the ideal treatment age for this study should be centered on the development of this region. In rodents, the peak development of the hippocampus occurs during the perinatal period. During this time, the hippocampus appears to be the most sensitive to hypoxia (Ikeda et al 2001) and drugs of abuse like methamphetamine (Williams et al 2003; Skelton et al 2007). The results of this study similarly suggest that the lack of Cr during peak levels of hippocampal development leads to cognitive deficits. Interestingly, conditioned fear behavior was not affected in either group of mice tested. Conditioned fear is thought to be based primarily in the amygdala (Butler et al 2017; Ressler and Maren 2018), which reaches its peak development around gestational day 12 in the mouse (Clancy et al 2007). The lack of a conditioned fear phenotype and the presence of deficits in hippocampally-based behaviors in the P5KO mice suggest that Cr may play an important role in the development of this brain region as well. Future studies could be designed to eliminate the *Slc6a8* gene in the developing amygdala and determine if the conditioned fear effect could be isolated.

The P5KO and the P60KO mice weighed less than their respective controls, leading to a possible concern that the smaller stature of these mice interferes with the interpretation of the cognitive data. The visible platform is frequently used to test for sensorimotor deficits (Vorhees and Williams 2006) and the P5KO mice perform this task slower than their WT-VEH counterparts. However, mice with a brain-specific *Slc6a8* deletion show visible platform deficits and do not have weight differences (Udobi et al 2018). In addition, the P5KO mice do show improvement over trials, suggesting that they are able to perform and learn the task. Finally, the P5KO mice do not show swim speed deficits during the hidden platform trials suggesting that the loss of Cr does not directly impact physical performance. Spatial learning and memory as well as novel object recognition deficits were observed in mice lacking *Slc6a8* in *Camk2a*-expressing neurons (Kurosawa et al 2012). While the primary focus of the Kurosawa study was on the therapeutic potential of cyclocreatine, the use of the Camk2a-Cre leads to a perinatal deletion of the *Slc6a8* gene (Casanova et al 2001), though only in excitatory neurons. Interestingly, the lack of a probe trial effect during the acquisition phase of the MWM was evident in both models. A probe trial effect was seen during reversal, showing that this extra testing phase is important to fully assess spatial learning and memory. As with the brain-specific *Slc6a8* knockout mice, the *Camk2a-Cre::Slc6a8*^*FLOX*^ mice did not have changes in body weight or swim speed, supporting the hypothesis that the deficits in the MWM are not due to motor changes in the mice. Cyclocreatine treatment during adulthood was able to rescue the cognitive deficits in the *Camk2a::Slc6a8*^*FLOX*^ mice. This would suggest that even though early Cr loss is required for deficits to manifest, the brain is still amenable to later treatment. It should be noted that the *Camk2a::Slc6a8*^*FLOX*^ mice may not be ideal for treatment due to an intact Slc6a8 at the blood brain barrier. Further studies using ubiquitous knockouts and restoration of the *Slc6a8* gene could provide a greater insight into this critical issue.

Increased respiration was observed in isolated mitochondria from *Slc6a8*^*−/y*^ mice, though this was likely due to an increase in total mitochondria as the tissues were normalized for total wet weight (Perna et al 2016). Increases in mitochondrial complex V and citrate synthase have been observed in *Agat*^*−/−*^ mice (Nabuurs et al 2013), further suggesting that changes in Cr content lead to mitochondrial dysfunction. In this study, no differences in stimulated respiration were observed in hippocampal synaptosomes isolated from P5 or P60 groups. This could suggest that the differences in mitochondrial content observed are localized to the cell body or other cell types such as astrocytes.

Attention deficit hyperactivity disorder is a common co-morbidity of CTD though it is not present in all patients (van de Kamp et al 2013). Accordingly, to date there has not been a consistent locomotor phenotype in *Slc6a8* mice. The *Slc6a8*^*−/y*^ mice showed an initial hypoactivity followed by similar levels to WT (Skelton et al 2011). Female heterozygous (Hautman et al 2014) and brain-specific knockouts were hyperactive (Udobi et al 2018). While P5KO mice were hyperactive during the dark phase compared with the WT-VEH mice, this was likely due to tamoxifen administration since all mice treated with tamoxifen from P5-10 were hyperactive. To our knowledge this is the first study that has shown that neonatal tamoxifen exposure leads to nocturnal hyperactivity. In a recent study, Mikelman et al show that tamoxifen interacts with the dopamine transporter (DAT) and can block amphetamine-stimulated dopamine release as well as blocking dopamine reuptake in rat synaptosomes (Mikelman et al 2018). While acute tamoxifen exposure did not change locomotor behavior, it was effective at blocking the locomotor-stimulating effects of amphetamine (Mikelman et al 2017; Mikelman et al 2018). It is possible that the neonatal DAT stimulation by tamoxifen caused the mild hyperactivity when the mice were tested as adults, though it is also possible that modulation of the estrogen receptor played a role. Together, these data add to the growing literature that behavioral studies employing tamoxifen, especially those looking at dopamine-modulated behaviors, should always include vehicle-treated controls. It should be noted that tamoxifen exposure did not lead to deficits in the cognitive tasks-showing that the loss of the *Slc6a8* was responsible for these deficits.

In conclusion, the findings of this paper show that Cr plays an important role in brain development in terms of cognition. Further, these data suggest that studies seeking to investigate possible treatments for CTD should consider starting treatment perinatally. This will allow for the maximum benefit of the compound as well as examining for any potential interactions between treatment and brain development.

**Table 1.**
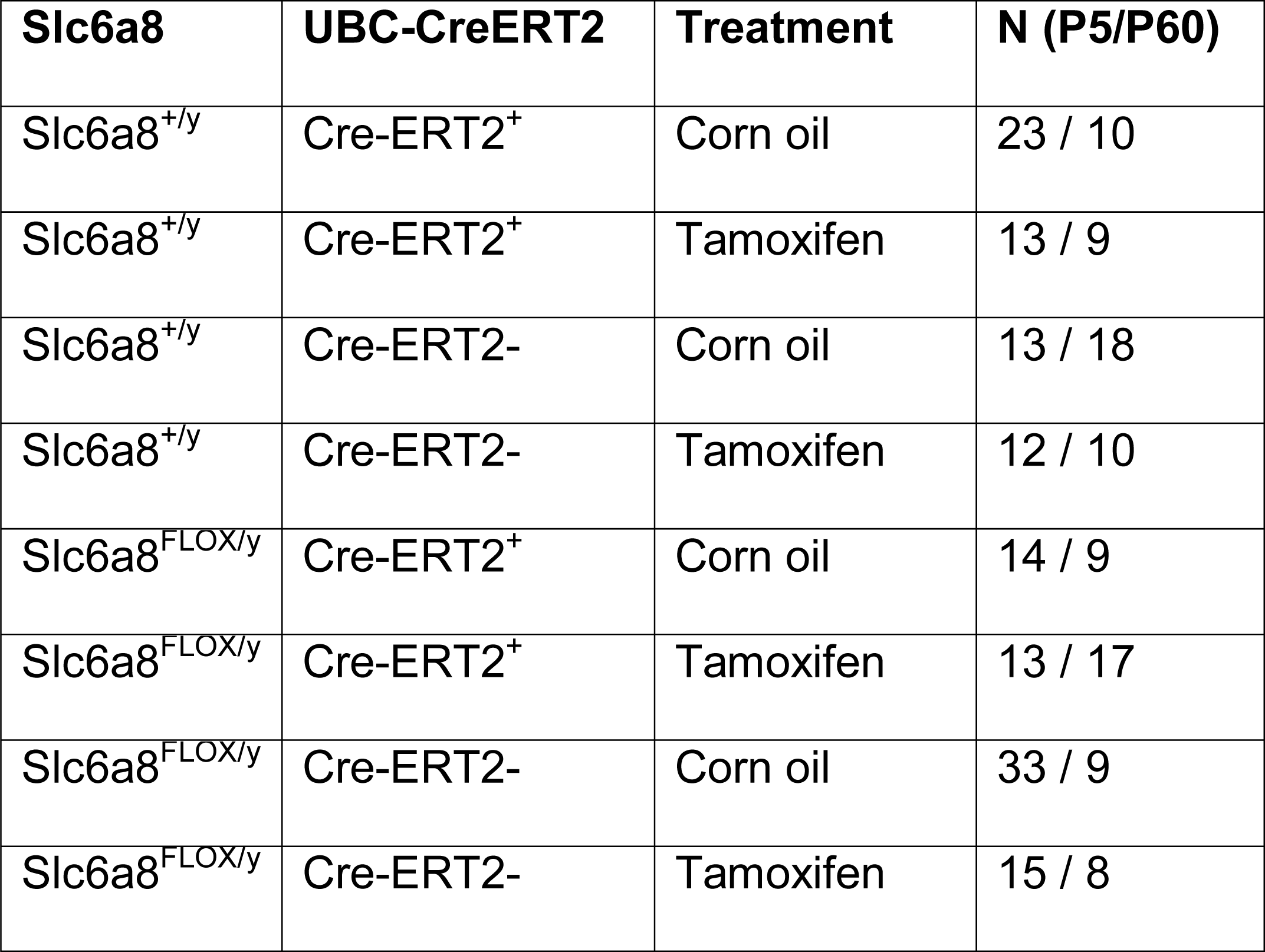
Subjects used.

**Table 2.** F Statistics for main effect of group

| Variable | Age | F statistic (nDF, dDF) | P value |
| --- | --- | --- | --- |
| Body Weight | P5 | F(1,175)=38.21 | <0.0001 |
|  | P60 | F(1,111)=167.23 | <0.0001 |
| MWM-Cued D1 | P5 | F(7,129)=2.83 | 0.009 |
|  | P60 | F(7,62)=1.29 | 0.2725 |
| MWM-Cued D2&D3 | P5 | F(7,126)=3.05 | 0.0054 |
|  | P60 | F(7, 61.9)=1.25 | 0.2925 |
| MWM-ACQ:<br>Latency | P5 | F(7,137)=3.97 | 0.0006 |
|  | P60 | F(7,97.1)=3.18 | 0.0045 |
| MWM-REV:<br>Latency | P5 | F(7,141)=1.83 | 0.0859 |
|  | P60 | F(7,96.9)=0.86 | 0.5432 |
| MWM-ACQ-Probe | P5 | F(7,137)=0.79 | 0.5964 |
|  | P60 | F(7,77)=1.06 | 0.3951 |
| MWM-REV-Probe | P5 | F(7,137)=1.72 | 0.1090 |
|  | P60 | F(7,77)=0.60 | 0.7555 |
| NOR | P5 | F(7,113)=2.29 | 0.032 |
|  | P60 | F(7,69)=0.87 | 0.5336 |
| CF-D2 | P5 | F(7,160)=0.78 | 0.6024 |
|  | P60 | F(7,86.1)=0.5 | 0.8291 |
| CF-D3-Tone | P5 | F(7,138)=1.47 | 0.1841 |
| | P60 | $F(7,68.9)=0.48$ | 0.8492 |
| Locomotor-Block 1 | P5 | $F(7,178)=1.02$ | 0.4182 |
| | P60 | $F(7,111)=1.99$ | 0.0627 |
| Locomotor-Block 2 | P5 | $F(7,459)=10.3$ | $<0.0001$ |
| | P60 | $F(7,299)=3.81$ | 0.0006 |
| Locomotor- Block 3 | P5 | $F(7,152)=3.19$ | 0.0035 |
| | P60 | $F(7,91)=0.51$ | 0.826 |

## Author contributions

Kenea Udobi and Matthew Skelton designed the research, analyzed the data, and drafted the manuscript. Nicholas Delcimmuto, Amanda N. Kokenge, Zuhair I. Abdulla, and Marla K. Perna performed the research.

## Acknowledgements

This work was supported by NIH grant HD080910, a CARE grant from the Association for Creatine Deficiencies, and internal support from the Division of Neurology (MRS). The authors thank Keila Miles for editorial support.

## Supplemental information

### Subjects

Mice were generated and maintained on a C57BL/6J background in a pathogen-free facility in microisolators. Female mice were not used since CTD is X-linked and primarily affects males. No more than one mouse per group was taken from a litter. All procedures on mice were performed at the CCRF vivarium which is fully accredited by AAALAC International and protocols were approved by the Institutional Animal Care and Use Committee. Lights were on a 14:10 light:dark cycle, room temperature was maintained at 22±1° C, and food (NIH-007) and filtered, sterilized water were available *ad libitum*. All institutional and national guidelines for the care and use of laboratory animals were followed.

### Cr determination by HPLC

A hemisphere was homogenized in 10 µL/mg tissue of 0.3N perchloric acid; homogenate was centrifuged at 4 °C for 10 min (15300g). Cr content was measured at 4 °C on a TSK gel ODS-80TM column (250 × 4.6 mm, 5 μm particle size, Tosoh, Tokyo, Japan) fitted with a guard column packed with C18 Nucleosil material. The mobile phase consisted of 50 mM NaH2PO4 (pH 2.15) and was degassed by vacuum filtration and sonication. A flow rate of 1 ml/min was used, and peaks were detected by absorbance at 210 nm. After each sample set (typically 20–40 injections of 90 μl each), the column was washed with 20% and 100% methanol.

### Real-time PCR

Total RNA was using TRIzol reagent (invitrogen). cDNA was synthesized from 0.5 ug total RNA using the iScript cDNA synthesis kit (Bio-rad, Hercules, California). All primers (Supplemental data) were designed for mouse gene sequence (Mus musculus taxid: 10090) using the web-based program Primer-BLAST (http://www.ncbi.nlm.nih.gov/tools/primer-blast/). Primers were obtained from Integrated DNA Technologies (IDT). Specific primer sequences are: *Slc6a8:* GCTTCCCCTACCTGTGCTAC and GGCATAGCCCAGACCTTTGAA; *Actb:* CCTGTGCTGCTCACCGAGGC and GCACAGTGTGGGTGACCCCG. qPCR was performed using an Applied Biosystems 7500 Real-Time PCR system with SYBR green mastermix containing ROX reference dye (KCQS01, Sigma-Aldrich). A 2-step QPCR was used to amplify mRNA; samples were denatured for 5 min at 95° C followed by 40 cycles of 95° C for 15 s and 65° C for 32 s.

### Morris Water Maze

The tank was 122 cm in diameter, white, and filled with room temperature (21 ± 1°C) water.

#### Visible Platform

During this phase, a 10 cm diameter platform with an orange ball mounted above the water on a brass rod was placed in one quadrant. Curtains were closed around the pool to obscure distal cues. On day 1, mice were given 6 trials with identical start and platform positions as training for the task. Mice were then given 2 trials per day for 2 days with the start and platform positions randomized.

#### Hidden platform

Hidden platform trials were conducted in 2 phases; each consisting of 4 trials per day for 4 days followed by a single probe trial without the platform on day 5. Platform diameters were 10 cm during acquisition and 7 cm during reversal (platform located in the opposite quadrant).

Mice were tested in a round-robin fashion with 8-10 mice completing trial 1 before trial 2 was started. The trial limit was 90 s and mice that reached the time limit were placed on the platform for the same amount of time as a mouse finding the platform (∼10 s).

### Novel object recognition

Mice were habituated to the arena (40 × 40 cm) for 2 days (10 min/day) followed by two days where they were placed in the arena with two identical objects (10 min/d) for 2 d to reduce neophobia (Podhorna and Brown 2002). Time exploring an object was defined as entry into a 2 cm zone around the object. Trials were recorded, and hand-scored by an experimenter blind to the groups. If a mouse failed to accumulate 30 s of total exploration time, the mouse was excluded from analysis. In the P5 group, 3 WT-VEH and 4 P5KO mice were excluded for failing to acquire the 30 s of observation time.

### Conditioned Fear

Each tone foot-shock pairing consisted of the 30 s tone accompanied during the last second by a scrambled footshock (82 dB, 2 kHz, 0.3 mA for 1 s) delivered through the floor. On day 1, mice were placed in a chamber for 3 min before exposure to 3 tone-footshock pairings. On day 2, mice were returned to the chamber with no tone or shock as a test of contextual fear. On day 3, mice were placed in the chamber with a novel floor for 6 min and exposed to the tone associated with the footshock for the final 3 min. The number of beam breaks was recorded using PAS software from San Diego instruments.

### Locomotor Behavior

Photobeam Activity System (PAS) – Open-Field (San Diego Instruments, San Diego, CA). Chambers were 40 cm (W) □ 40 cm (D) □ 38 cm (H) with 16 LED-photodetector beams in X and Y planes was used to measure locomotor activity. Photocells were spaced 2.5 cm apart. The overhead lighting is controlled with a timer that mimics the phases used in the housing suite with lights on at 0600 and off at 2000.

### Statistics

Data were analyzed using mixed-model ANOVA (Proc Mixed, SAS, Cary, NC) followed by post-hoc Dunnett’s test with all groups compared with the WT-VEH. Kenward-Roger degrees of freedom were used. For the MWM, conditioned fear, and locomotor behavior, trial or day was included as a repeated measure.

---

**Supplemental Figure 1.**
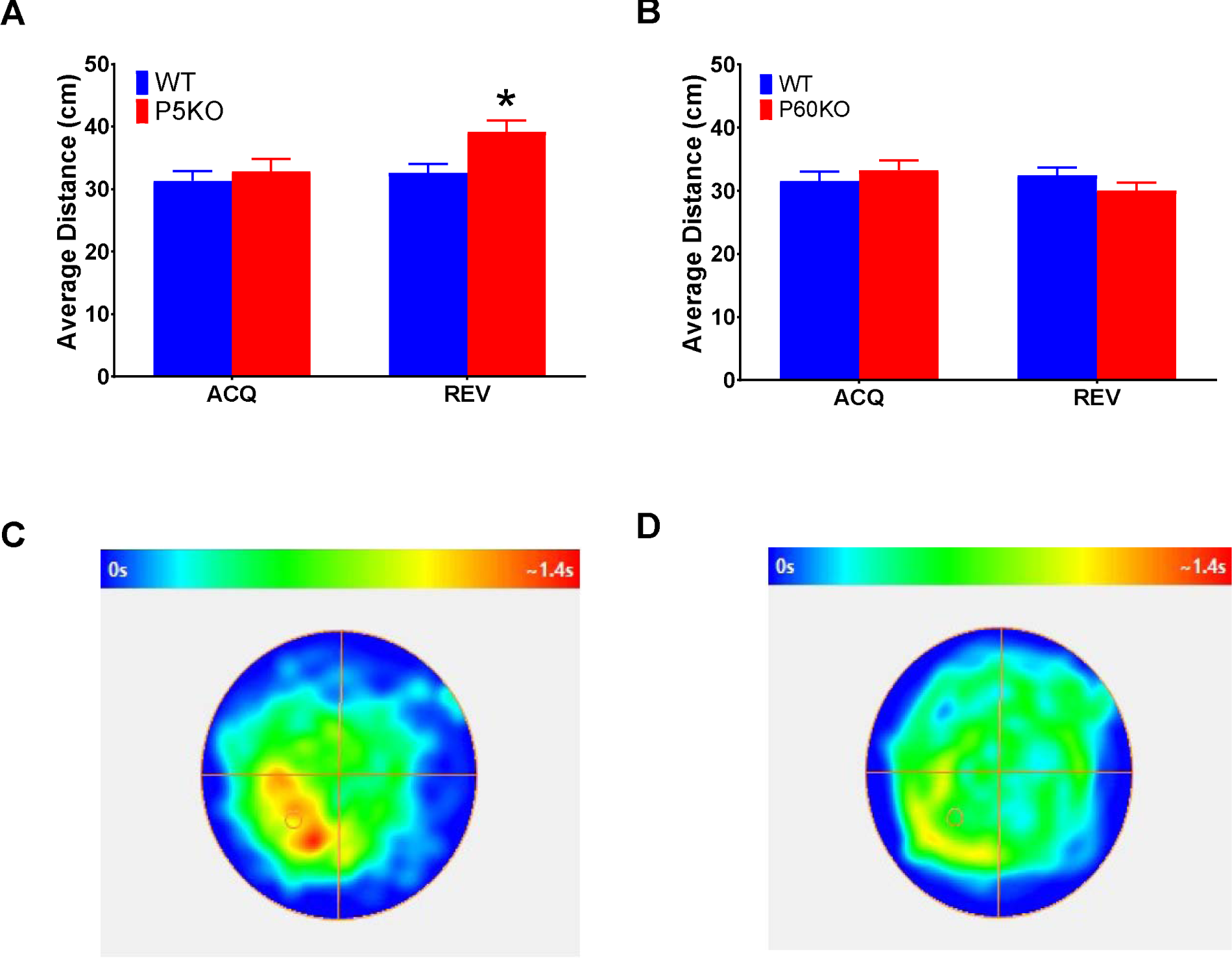
Spatial memory deficit in P5KO mice. (A) P5KO mice had an increased average distance from the target platform during the reversal (REV) probe trial. No difference in average distance was observed during acquisition (ACQ). (B) P60KO mice performed similarly to WT-VEH during acquisition and reversal probe trials. When looking at the composite heat-map data from the reversal trial, the WT-VEH mice (C) show more time near the platform site compared with P5KO mice (D). Data are LSMEAN±SEM, *p<0.05. N=13-18/group

**Supplemental Figure 2.**
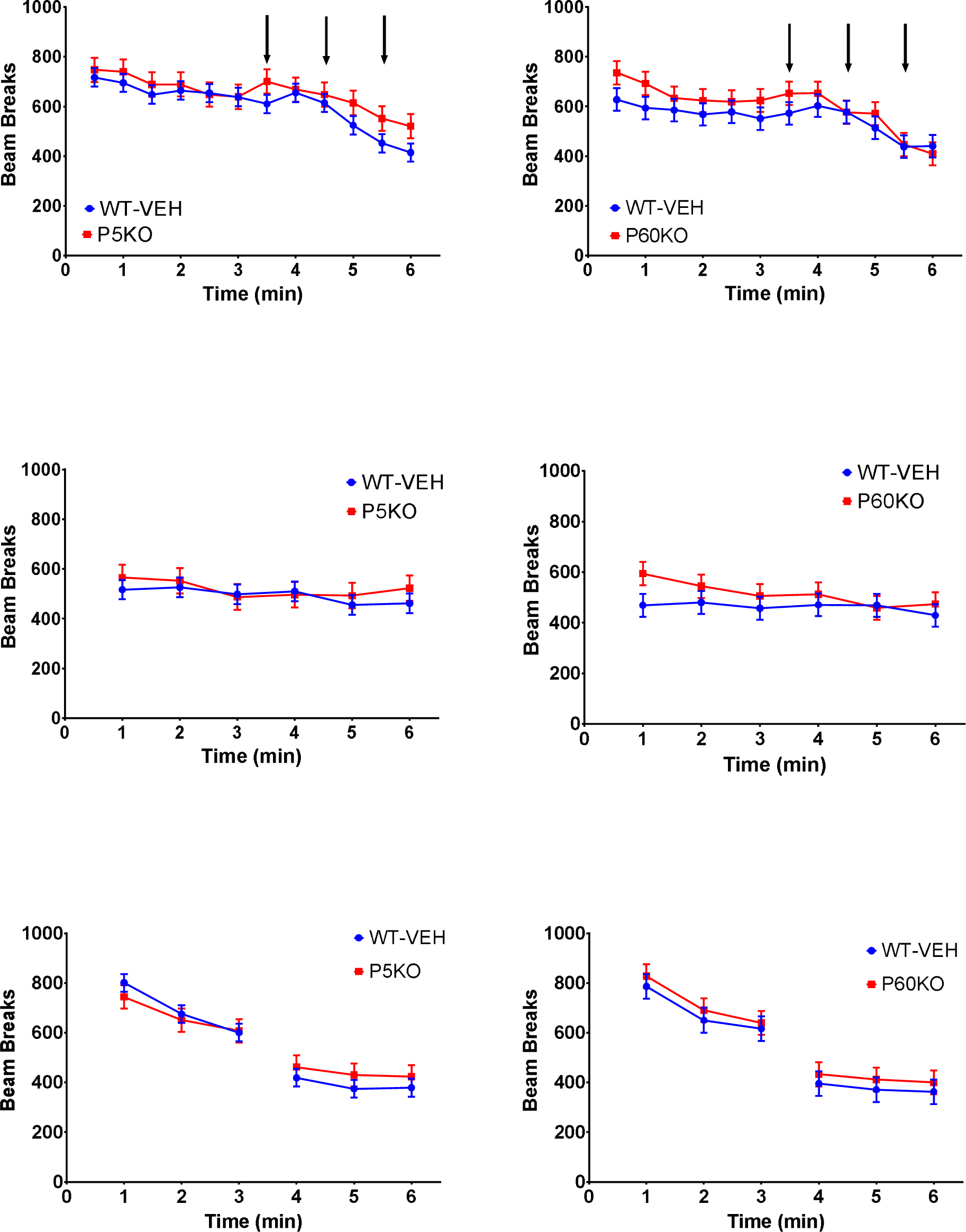
Conditional *Slc6a8* knockout does not affect fear memory. *Top:* There were no differences in activity levels before, during (arrows), or following the tone-footshock conditioning on day 1. *Middle:* Both treatment ages performed similarly to their corresponding WT-VEH mice when tested for contextual fear. The activity levels during this task are similar to the levels following the tone-footshock pairing on the previous day, suggesting the mice associated the arena with the footshock. *Bottom:* No differences in activity levels before or during tone presentation in either P5KO or P60KO mice. Note the reduction in activity corresponding with a successful pairing of the tone to the footshock. Data are LSMEAN±SEM. N=13-18/group.

**Supplemental Figure 3.**
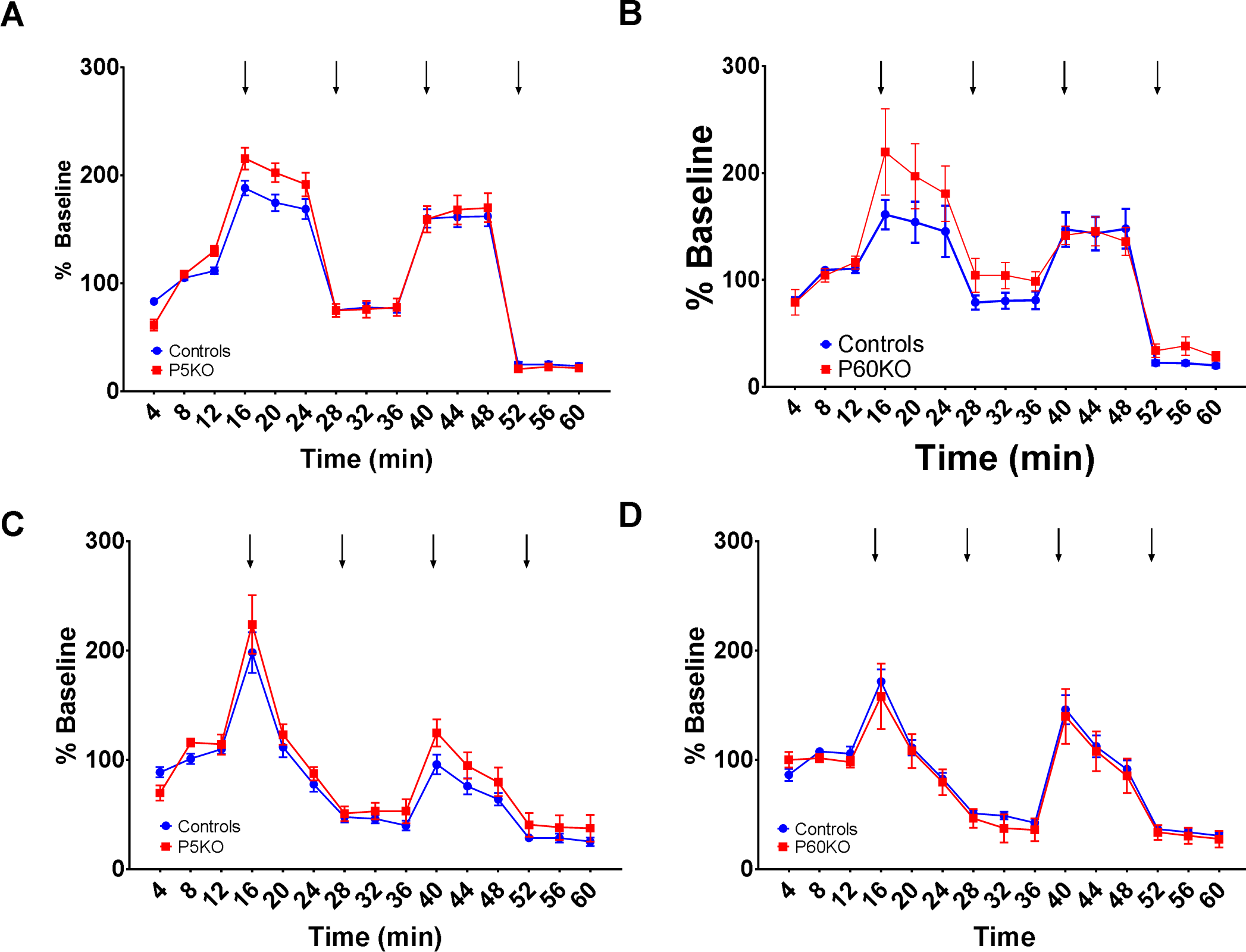
Stimulated respiration in hippocampal synaptosomes. Panels A and B are synaptosomes incubated with rotenone and succinate. Panels C and D are synaptosomes incubated with malate and pyruvate. Arrows represent injections of (in order): a) ADP; b) oligomycin; c) FCCP; d) Antimycin A.

